# Continuous Fish Muscle Cell Line with Capacity for Myogenic and Adipogenic-like Phenotypes

**DOI:** 10.1101/2022.08.22.504874

**Authors:** Michael K Saad, John SK Yuen, Connor M Joyce, Xinxin Li, Taehwan Lim, Talia L Wolfson, Justin Wu, Jason Laird, Sanjana Vissapragada, Olivia P Calkins, Adham Ali, David L Kaplan

## Abstract

Cell-cultivated fish offers the potential for a more ethical, sustainable, and safe seafood system. However, fish cell culture is relatively understudied in comparison to mammalian cells. Here, we established and characterized a continuous Atlantic mackerel (*Scomber scombrus*) skeletal muscle cell line (“Mack” cells). The cells were isolated from muscle biopsies of fresh-caught fish, with separate isolations performed from two distinct fish. Mack1 cells (cells from the first isolation) were cultured for over a year and subcultured over 130 times. The cells proliferated at initial doubling times of 63.9 hr (± 19.1 SD). After a spontaneous immortalization crisis from passages 37-43, the cells proliferated at doubling times of 24.3 hr (± 4.91 SD). A muscle phenotype was confirmed through characterization of muscle stemness and differentiation via paired-box protein 7 (PAX7) and myosin heavy chain (MHC) immunostaining, respectively. An adipocyte-like phenotype was also demonstrated for the cells through lipid accumulation, confirmed via Oil Red O (ORO) staining and quantification of neutral lipids. New qPCR primers (*HPRT*, *PAX3B*, *MYOD1*, *MYOG*, *TNNT3A*, and *PPARG*) were tailored to the mackerel genome and used to characterize mackerel cell genotypes. This work provides the first spontaneously immortalized fish muscle cell line for research, ideally serving as a reference for subsequent investigation.

## Introduction

Cell culture research on aquatic and marine species is relatively understudied when compared to mammalian cell culture research, with zebrafish as an exception. In cellular agriculture, which is the production of animal-sourced foods via cell culture, immortalized cells are essential for consistency during large scale production. Most *in vitro* cell culture studies for future foods have focused on livestock animals, such as beef cattle^1–4^. However, due to concerns of overfishing and fish health in natural environments due to heavy metals, pesticides, and plastics, as well as the negative ecological impact of aquaculture, *cellular aquaculture*, or cell-cultivated fish, is an emerging yet understudied realm of cellular agriculture. Currently, fish support 15% of the global demand for animal protein; coupled with a growing population on the planet, the demand for fish is projected to rise further^5^. Current aquaculture and marine fish capture processes have significant drawbacks in terms of food safety, depletion of fish stocks, and the environment. Overfishing is deleterious to marine ecosystems and is depleting numerous fish species to the point of extinction^6^. In terms of food safety, foodborne diseases, plastics accumulation, and heavy metal contamination are major issues, while environmental concerns connected to climate change are damaging marine diversity^7,8^. Cellular aquaculture has the potential to alleviate fish sourcing concerns, with a focus on production efficiency, scaling, and controlled cultivation environments to address food security concerns. However, to date there are no available fish skeletal muscle cell lines to produce cell-cultured fish meat, or to use as a model for research, like immortalized C2C12 mouse myoblasts for mammalian muscle cell culture.

Nine fish cell lines, from *Acipenser transmontanus* (white sturgeon), *Caranx melampygus* (bluefin trevally), *Lates calcarifer* (barramundi), *Danio rerio* (zebrafish), *Carassius auratus* (grass goldfish), *Channa striata* (snakehead murrel), *Labeo catla* (Indian catla), and *Wallago attu* (helicopter catfish), have been isolated from fish muscle^9,10^; however, none of the cell lines are characterized as muscle cells, as they lack myogenic potential. While fish muscle cells from species such as *Danio rerio* (zebrafish), *Salmo salar* (Atlantic salmon), *Carassius auratus* (grass goldfish), and *Oncorhynchus mykiss* (rainbow trout) have been isolated, their applicability to cellular aquaculture has been limited^11–14^. These limitations stem from some of these species not being commonly consumed (e.g., zebrafish, goldfish), the cell population not demonstrating muscle function (e.g., the aforementioned cell lines, goldfish), or their limited growth potential (salmon, trout) as the cells were not demonstrated in long-term culture. For cellular aquaculture, a cell line with applicability to food through demonstration of myogenic or adipogenic functions in long-term culture would provide a reference for research as well as applicability for food production processes. In particular, a cell line from a commonly consumed fish species has direct benefits for potential commercialization, as consumers would likely favor *in vitro* fish from a recognized source.

Atlantic mackerel (*Scomber scombrus*) is an understudied fish species, with few *in vivo* research studies and no *in vitro* cell studies published. Mackerel is a popular fish for consumption, and the oils from this fish are desired for health benefits due to their high percentage of omega-3 fatty acids^15^. However, mackerel populations have significantly declined in recent years, with the Monterey Bay Aquarium’s Seafood Watch, a platform for identifying environmentally sustainable seafood, recommending avoiding consumption of the fish due to its depleted stocks and exploitative overfishing^16^. Additionally, in 2004, the US Food and Drug Administration (FDA) advised against eating mackerel due to high levels of mercury^17^. As an alternative to fished mackerel, the isolation and propagation of mackerel cells (muscle, fat) towards cell-cultivated mackerel tissue offers an option to address the above concerns.

The objective of this work was to isolate and characterize mackerel muscle cells towards a continuous cell line via spontaneous immortalization. First, since mackerel is wild-caught and confirmation of species identity is a high priority in the regulation of fish and their labeling^18^, we confirmed the cells’ identity as *S. scombrus* through PCR on cell genomic DNA and subsequent Sanger sequencing. After establishment of a continuous culture, past a spontaneous immortalization crisis event, we confirmed myogenicity through immunocytochemistry for paired-box protein 7 (PAX7) in proliferating cells, a marker for muscle cell stemness, and myosin heavy chain (MHC) in differentiated cells. As mackerel is an oily fish, we also demonstrated the cells’ ability for lipid accumulation towards adipogenic-like outcomes. We investigated cell culture optimization, determining optimal seeding and differentiation methods while screening various extracellular matrix (ECM) surface coatings. Finally, we initiated the development of research tools, overcoming difficulties due to the lack of annotations on the *S. scombrus* genome and including the design of primers for future gene expression studies and research needs.

## Methods

### Primary mackerel satellite cell isolation and maintenance

Isolations were performed on two separate fish, and cells from the first and second isolations are referred to as Mack1 and Mack2, respectively. Primary mackerel satellite cells were isolated enzymatically using collagenase. Briefly, wild-caught, deceased Atlantic mackerel (*S. scombrus*) was transported to the lab on ice. The isolations generally began ^~^5 hr *post-mortem*. The whole fish was first dipped in 70% ethanol for 10 seconds before being transferred into the tissue culture hood. Then, ^~^5 g of epaxial white muscle tissue was excised then sterilized in 70% ethanol and rinsed in Dulbecco’s phosphate-buffered saline (DPBS, ThermoFisher Scientific #14190144, Waltham, MA, USA). The muscle tissue was minced into a paste and digested in DPBS containing 0.1% collagenase, type II (Worthington Biochemical #LS004176, Lakewood, NJ, USA) for 1 hr at room temperature, with trituration every 15 min. The digested tissue was strained through 100, 70, and 40 μm strainers. The strained cell suspension was centrifuged at 300 x g to pellet the cells and then resuspended in 10 mL of growth medium, comprised of Leibovitz’s L-15 Medium (ThermoFisher Scientific #11415064) with 20% fetal bovine serum (FBS; ThermoFisher Scientific #26140079), 1 ng/mL recombinant human fibroblast growth factor (FGF-basic 154 a.a., PeproTech #100-18B, Cranbury, NJ, USA), 20 mM HEPES pH 7.4 (Sigma Aldrich #H4034, St. Louis, MO, USA), 1% Antibiotic-Antimycotic (ThermoFisher Scientific #1540062), and 10 μg/mL gentamicin (Sigma Aldrich #G1397). The cells in growth medium were then counted using a NucleoCounter NC-200™ automated cell counter (Chemometec, Allerod, Denmark) and seeded at ^~^100,000 cells/cm^2^ into tissue culture-treated flasks. After 48 hr of incubation at 27°C without CO2, the plated suspensions were transferred to flasks coated with 0.25 ug/cm^2^ iMatrix recombinant laminin-511-E8 (Iwai North America #N892021, San Carlos, CA, USA). The cells were not disturbed for 5 days to allow for adhesion. The cells were then fed until they reached ^~^70% confluency, about 10 days after the isolation. Cells were maintained by passaging at ^~^70% confluency and seeding between 2,000 – 8,000 cells/cm^2^ or stored by freezing in growth medium with 10% dimethyl sulfoxide (DMSO, Sigma Aldrich #D2650). During early passages, 0.25% trypsin-EDTA (ThermoFisher Scientific #25200056) was used to liberate the cells from the surfaces. 0.05% trypsin-EDTA (ThermoFisher Scientific #25300054) was used after a spontaneous immortalization crisis event, characterized by high incidence of senescent cell morphology in the culture, at around passage 37 (P37).

### PCR identification

To confirm the identity of the cells from *S. scombrus*, genomic DNA from *in vitro* culture (Mack1 P4 (n = 2), Mack1 P55 (n = 1), and Mack2 P3 (n = 1)) was purified using GeneJET Genomic DNA Purification Kits (ThermoFisher Scientific #K0721). In an adapted method from Aranishi^19^, PCR was performed with Q5^®^ High-Fidelity 2X Master Mix (New England Biolabs (NEB) #M0492, Ipswich, MA, USA), with primers designed to target fish universal 5S rDNA (5S primers) and *Scomber* genus-specific sequences of the 5S rDNA (Saba primers). The 5S forward and reverse primers used were 5S21F (5’-TACGC CCGAT CTCGT CCGAT C-3’) and 5S21R (5’-CAGGC TGGTA TGGCC GTAAG C-3’), respectively. The Saba forward and reverse primers used were Saba-18F (5’-GGGCG CTGTT GCTCC ATC-3’) and Saba-20R (5’-ATGCT GTGAC ACCAC TGACA-3). To confirm successful PCR, the amplification products were run on a 2% agarose gel and imaged on the G:BOX Chemi XR5 (Syngene, Bangalore, India). To confirm species identity, the 5S and Saba PCR products were purified using Monarch^®^ PCR & DNA Cleanup Kit (5 μg) (NEB, # T1030). The purified DNA sequence was then analyzed through Sanger sequencing (GENEWIZ from Azenta, South Plainfield, NJ), using the forward primer for amplification. The sequences were aligned to known 5S rDNA sequences for *S. colias, S. scombrus, S. australasicus*, and *S. japonicus* (NCBI GenBank Accession Numbers AB245986.1, AB246033.1, AB246025.1, and AB246016.1, respectively) with Geneious Prime (Biomatters, Inc., San Diego, CA, USA) nucleotide alignment.

### Doubling time and cell diameter measurements

To track doubling time during cell culture, the dates/times of subculture and total viable cell numbers, as reported by the NC-200 automated cell counter, were recorded. The growth rate and doubling time were calculated using eqs. 1 and 2:

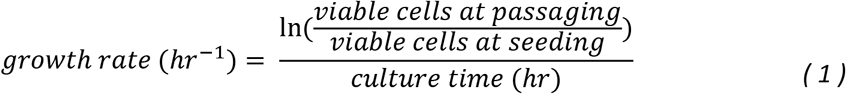

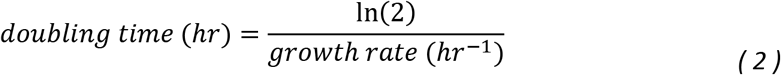

Cell diameters were obtained with the NC-200 automated cell counter.

### Immunoctyochemistry

To validate myogenic potential, proliferating mackerel satellite cells were stained for paired-box protein 7 (PAX7), a marker of satellite cell identity. Cells were fixed with 4% paraformaldehyde (ThermoFisher Scientific #AAJ61899AK) for 30 min, washed in DPBS with 0.1% Tween-20 (PBST, Sigma Aldrich #P1379), permeabilized for 10 min using 0.1% Triton-X (Sigma Aldrich #T8787), blocked for 30 min using 1X blocking buffer (Abcam #ab1265875, Cambridge, UK) in DPBS, and washed with PBST. Primary PAX7 antibody (ThermoFisher Scientific #PA5-68506) was diluted 1:100 in 1X blocking buffer, added to cells, and incubated overnight at 4°C. Cells were then washed with PBST, blocked for 30 min using 1X blocking buffer, and incubated for 1 hr with secondary antibody (ThermoFisher Scientific, #A-11072) and Phalloidin-iFluor 488 Reagent (Abcam #ab176753), both diluted 1:1000 in 1X blocking buffer. After washing with PBST, the cell nuclei were stained with 1 μg/mL 4’,6-diamidino-2-phenylindole (DAPI, ThermoFisher Scientific #62248) in DPBS for 15 min at room temperature. Imaging was performed with a fluorescent widefield microscope (KEYENCE, BZ-X700, Osaka, Japan).

To verify myogenicity, cells were cultured to 90-100% confluency in growth medium, after which cell feeding was ceased (termed “serum starvation”). Following 10 days of differentiation, the cells were fixed, stained, and imaged as previously described, using primary antibodies for myosin heavy chain (Developmental Studies Hybridoma Bank #MF-20, Iowa City, IA, USA; 4 μg/mL), appropriate secondary antibodies (ThermoFisher Scientific #A-11001, 1:1000), Phalloidin, and DAPI.

### Image analysis of immunostaining

To quantify the percentage of nuclei expressing either PAX7 or MHC, the BZ-X800 Analyzer software (KEYENCE, Osaka, Japan) was used. Overlay images with cell nuclei (DAPI) and the immunostained target of interest were analyzed using the “Hybrid Cell Count” software for single extraction of fluorescence images. Results from the analysis were then run through a custom Python script to remove any replicate counts within the same nuclei. For the PAX7 expression comparison over multiple passages, the samples were first blinded and then passed through the analysis.

Muscle fiber diameter analysis was performed using ImageJ. First, the scale bar in the image was used to set the measuring scale in the program. The line tool was then used to outline the diameter of the muscle fiber that needed to be measured. Then, the measuring command was used to record the muscle fiber diameter. The measurements were taken randomly, with n = 20 for each image.

### Lipid accumulation

For lipid accumulation towards an adipogenic-like phenotype, cells were grown to 100% confluency and then fed 3 times a week with adipogenic medium (growth medium with 10 μg/mL insulin (Sigma Aldrich #I0516), 0.5 mM 3-isobutyl-1-methylxanthine (IBMX, Sigma Aldrich #I5879), 0.25 μM dexamethasone (Sigma Aldrich #D4902), and 10 μL/mL lipid mixture (Sigma Aldrich #L5146)). To visualize neutral lipids after 7 and 14 days of lipid accumulation, cells were fixed then stained with Oil Red O (ORO, Sigma Aldrich #O0625) and imaged via fluorescence microscopy. Briefly, a 3 mg/mL ORO stock was prepared by dissolving ORO powder in 100% 2-proponal (VWR #BDH2032-1GLP, Radnor, PA, USA). That stock solution was then diluted to a 1.8 mg/mL working solution in UltraPure™ Distilled Water (ThermoFisher #10977015), incubated at room temperature for 5 min, and then filtered through a 0.22 μm syringe filter. After fixation, the cells were incubated with ORO working solution for 10 min, washed with water, and then imaged. To quantify lipid accumulation, ORO was extracted from the cells with 2-propanol, and absorbance at 510 nm was measured on a Synergy H1 microplate reader (BioTek Instruments, Winooski, VT, USA).

### Lipidomics

Cells accumulated lipids for 14 days as described above, then harvested and the lipids extracted via methyl tert-butyl ether (MTBE, Alfa Aesar #41839AK, Haverhill, MA, USA) extraction as previously described^20,21^. Briefly, cells were grown in 3 x T75 flasks, rinsed with DPBS, and harvested with a cell scraper (Fisher Scientific #NC1890484, Waltham, MA, USA). Suspended cells were vortexed with methanol (ThermoFisher Scientific #BPA4544) before adding MTBE and incubating for 1 hr. Distilled water was added, then samples were vortexed and centrifuged at 10,000 x g for 5 min. The upper organic phase was removed, and MTBE extraction was repeated. Double-extracted MTBE phases were dried under liquid nitrogen and stored at −20°C.

Untargeted lipidomics profiling (n = 3) was performed via high-resolution liquid chromatography with tandem mass spectrometry (Thermo Scientific QExactive Plus/HF Orbitrap HR-LC-MS/MS) at the Mass Spectrometry Core at Beth Israel Deaconess Medical Center in Boston, Massachusetts^20^. Data processing was performed in Python (v 3.8.5). The code and preprocessed dataset are available in GitHub at the following URL: https://github.com/mksaad28/Saad_et_al_Mack. Python’s pandas (v 1.2.3) was used to organize the data for manipulation. The lipids were binned according to their classification (e.g., phospholipids, triglycerides). Within each classification, the lipid peak areas of the associated fatty acids were summed to determine total fatty acid area, and the fatty acids were sorted as saturated (SFAs), monounsaturated (MUFAs), or polyunsaturated (PUFAs) based on composition.

### Cell growth at different seeding densities

To determine the minimum seeding density and maximum cell confluency of mackerel cells, the viability of mackerel cells was measured using the CyQUANT™ Cell Proliferation Assay kit (ThermoFisher Scientific #C7026) according to the manufacturer’s instructions. Cells were seeded in 96-well plates at different seeding densities (1,000, 2,000, 3,000, 4,000, 5,000, 6,000, 8,000, and 10,000 cells/cm^2^) in triplicate for each time point and cultured at 27°C. The culture medium was replaced every other day. At each time point (days 1, 3, and 5), medium was aspirated, and the plate was stored at −80°C. After thawing and a 5-min incubation with CyQUANT™ working solution, fluorescence was measured at an excitation of 480 nm and emission of 520 nm using a Synergy H1 microplate reader. Cell number was calculated using a standard curve of cells seeded at a known density.

### Surface coatings

To assess ECM efficacy in optimizing cell growth, muscle differentiation, and adipogenic-like differentiation, different 2D surface coatings were studied. The following proteins and concentrations were used: no coating control; pre-coated collagen, type IV (Corning #354233, Corning, NY, USA) at 5 μg/cm^2^; pre-coated fibronectin bovine protein (ThermoFisher Scientific #33010018) at 10 μg/cm^2^; pre-coated gelatin (ATCC #PCS-999-027, Manassas, VA, USA) at 0.1 mg/cm^2^; laminin-511-E8 at 0.25 μg/cm^2^, added to the media; and pre-coated vitronectin recombinant human protein (ThermoFisher Scientific #A14700) at 0.5 μg/cm^2^.

Mack cells were seeded on the different surfaces at 6,000 cells/cm^2^ in experimental triplicate. The proliferation of mackerel cells was monitored at 1-, 3-, and 5-days post-seeding using FluoReporter™ Blue Fluorometric dsDNA Quantitation Kit (ThermoFisher Scientific #F2962) according to the manufacturer’s instructions. For FluoReporter™, fluorescence readings were performed on a Synergy H1 microplate reader using excitation and emission filters centered at 360 and 460 nm, respectively. Mackerel muscle differentiation was encouraged with cessation of feeding (serum starvation) and evaluated for MHC expression with MF-20 (immunostaining), as described earlier. Mackerel adipogenic-like differentiation was performed using adipogenic medium and evaluated with ORO quantification, as described earlier.

### Muscle differentiation method screen

To screen different muscle differentiation regimes, six different methods were tested: cessation of feeding (serum starvation); medium containing 2% FBS (reduced serum); a differentiation medium (reduced serum + additives) adapted from Messmer et al.^3^ – reduced serum with insulin (Sigma Aldrich #I0516), 1-oleoyl lysophosphatidic acid (LPA) (Fisher Scientific #NC9401387), and transferrin (InVitria #777TRF029, Aurora, CO, USA); reduced serum medium with insulin-like growth factor 1 (IGF-1; Shenandoah Biotechnology #100-34, Warminster, PA, USA); reduced serum medium + additives with IGF-1; and a medium with an extracellular signal-regulated kinase inhibitor (ERKi, MilliPore Sigma #1049738-54-6, Burlington, MA, USA), adapted from Eigler et al.^2^ More details on the media used are provided in Supplementary Table 1.

Mack1 cells were seeded with laminin-511-E8 in experimental triplicate and, upon reaching 100% confluency, were fed with the different regimes listed above. After 10 days in culture, cells were fixed, stained with DAPI and MF-20, and images analyzed for the percent of MHC+ nuclei, as described earlier.

### RT-qPCR

Primers for RT-qPCR were designed using the National Center for Biotechnology information (NCBI) Primer-BLAST tool. Primers were designed for the following southern bluefin tuna (*Thunnus maccoyii*) genes: hypoxanthine-guanine phosphoribosyltransferase (*HPRT*, a housekeeping gene), paired box 3b (*PAX3B*), myogenic differentiation 1 (*MYOD1*), myogenin (*MYOG*), (troponin T type 3a (*TNNT3A*), and peroxisome proliferator-activated receptor gamma (*PPARG*). For subsequent iterations of primer design, a *S. scombrus* transcriptome (NCBI SRX2255766) was aligned with a *T. maccoyii* annotated genome (NBCI RefSeq GCF_910596095.1) with STAR aligner (v.2.7.0a). Primer designs were then manually altered to better match the *S. scombrus* sequences.

For RT-qPCR, experimental triplicate cultures of Mack1 cells were prepared for four conditions: proliferation, muscle differentiation with serum starvation; muscle differentiation with reduced serum + additives (serum reduction); and lipid accumulation (adipogenic-like). For RNA extraction, cell lysates were harvested using the RNeasy Mini kit (Qiagen #74104, Germantown, MD, USA), according to the manufacturer’s instructions. RNA concentrations were determined by Nanadrop™ 2000 spectrophotometer (ThermoFisher Scientific #ND-2000) at 260 nm. After RNA concentrations were normalized to the concentration of the least concentrated sample, RNA was reverse-transcribed to cDNA using the iScript™ cDNA Synthesis kit (Bio-Rad #1708890, Hercules, CA, USA), according to the manufacturer’s instructions.

To screen for successful primers, PCR was performed on cDNA using Q5® High-Fidelity 2X Master Mix, and PCR products run on 2% agarose gels for visual confirmation of efficacy and specificity. After screening confirmation, RT–qPCR was performed with technical triplicates for each experimental replicate using SsoAdvanced Universal SYBR® Green Supermix (Bio-Rad #1725270) with primer pairs shown in Supplementary Table 2. Using the cycle threshold (Ct) values, ΔCt was calculated for each sample and gene of interest by subtracting the Ct of the housekeeping *HPRT* gene for each sample. ΔΔCt was then calculated by subtracting the proliferation ΔCt from the sample ΔCt for each gene. When gene expression was not detected for proliferation samples (as was observed in some replicates of *TNNT3A*), the average of the run’s other replicates for that gene was used. Fold induction over proliferation was calculated as 2^-ΔΔCt^.

### Mycoplasma Testing

Cell cultures were monitored for the presence of mycoplasma contamination using the MycoAlert® Mycoplasma Detection Kit (Lonza #LT07-318, Basel, Switzerland) according to the manufacturer’s instructions. The MycoAlert® Assay is a selective biochemical test that evaluates mycoplasma enzymatic activity by measuring conversion of adenosine diphosphate (ADP) to adenosine triphosphate (ATP). Spent medium was collected following > 24-hr incubation in cell culture then centrifuged at 200 x g for 5 min, and the supernatant was transferred into white-walled opaque bottom 96-well plate (Griener Bio-One #655083, Kremsmünster, Austria). Luminescence was measured using a Synergy H1 microplate reader microplate reader. Samples were incubated with MycoAlert™ Reagent for 5 min, and luminescence was measured (Reading A). Samples were then incubated with MycoAlert™ Substrate for 10 min, and luminescence was measured again (Reading B). The luminescence ratio of Reading B/Reading A was used to determine the presence of mycoplasma contamination with ratio < 0.9 interpreted as negative for mycoplasma contamination. Samples were tested in technical triplicate, and assay performance was validated using negative and positive controls (MycoAlert™ Assay Control Set, Lonza #LT07-518).

### Statistical analyses

Statistical analyses were performed with GraphPad Prism (v9.3.1, San Diego, CA, USA). For comparing means between two samples, unpaired t-tests were performed, as indicated in figure legends. For comparison of more than two samples, either one-way or two-way Analysis of Variance (ANOVA) was performed, as indicated in figure legends, with multiple comparisons performed via Tukey’s Honest Significant Difference (HSD) post-hoc test. P values < 0.05 were treated as significant. All errors are given as ± standard deviation, unless otherwise indicated. For RT-qPCR data, outliers were removed with JMP (v16.0.0), using the Univariate Quantile Range Outliers command, with the tail quantile = 0.3 and Q = 3.

## Results

### PCR confirmation of *S. scombrus* identity

To confirm the identity of the isolated cells as Atlantic mackerel, an adapted method from Aranishi^19^ was employed. PCR on cell genomic DNA was run with two sets of primers, fish universal 5S primers and *Scomber* genus-specific Saba primers (Fig. 1A). The amplification products matched the expected size of 486 base pairs (bp) and 311 bp for 5S and Saba reactions, respectively. To investigate the species specificity, PCR products were sequenced via Sanger sequencing. The sequencing results were then compared by alignment to four scombrid species, *S. colias* (Atlantic chub mackerel), *S. scombrus* (Atlantic mackerel), *S. australasicus* (Chub mackerel), and *S. japonicus* (Blue mackerel) (Fig. 1B). As *S. scombrus* exhibited 97.7% sequence identity to the consensus mackerel sequence (the consensus sequence from alignment of all the PCR product sequences), the identity of the cells was confirmed as Atlantic mackerel. While *S. scombrus* did exhibit slight deviations from Aranishi’s sequencing (e.g., nucleotide mismatch in position 129), the other *Scomber* species exhibited large insertions or deletions when compared against the DNA sequence of the cells (Fig. 1C).

**Figure 1:**
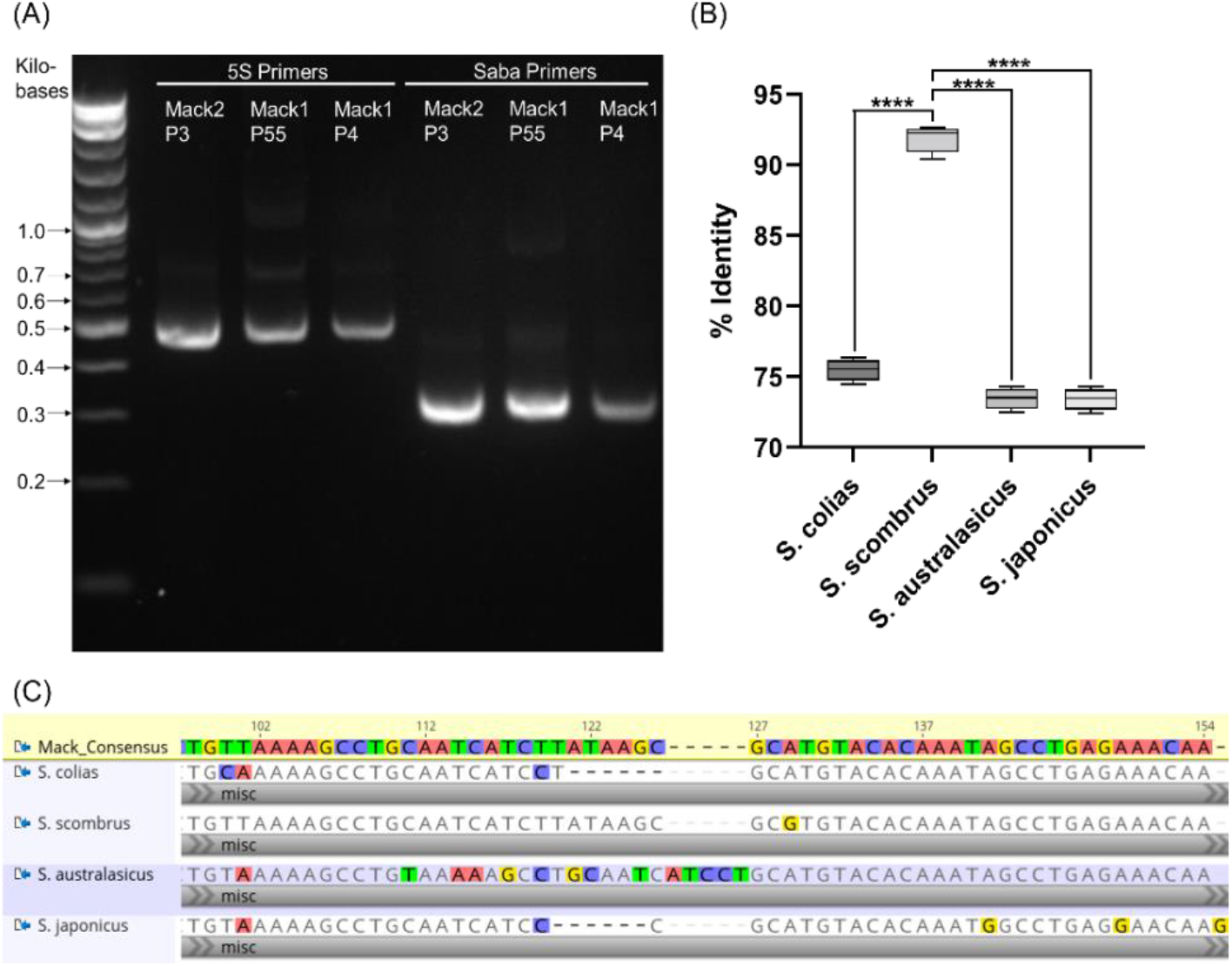
PCR confirmation of *S. scombrus* identity. (A) PCR amplification of Mack cell genomic DNA (Mack2 P3, Mack1 P55, Mack1 P4) with fish universal 5S primers and *Scomber*-specific Saba primers, with apparent band size matching expected size of 486 bp and 311 bp, respectively. (B) Comparison between *Scomber* species of Geneious Prime nucleotide alignment identity percentages of Sanger sequencing from Saba PCR reaction. Whiskers indicate min to max, n = 4; statistical significance calculated by one-way ANOVA with multiple comparisons between the control *S. scombrus* and the other species and indicated by asterisks, in which p < 0.0001 (****). (C) Section of Geneious Prime alignment of the consensus mackerel nucleotide sequence (n = 4) and the four *Scomber* species.

### Establishment of continuous cell culture through spontaneous immortalization crisis

The primary muscle cells were maintained in culture continuously by subculturing. To establish a cell line without genetic modifications, the cell culture proceeded through a crisis of spontaneous immortalization transformation, as previously described by Li *et al^12^*. Li *et al*. characterized the spontaneous immortalization crisis by a large-scale apoptosis/senescence, resulting in sporadic clusters of quiescent cells that then grow out of quiescence to reach more confluent areas^12^. From passages 37 to 43, the mackerel cells underwent a similar crisis, marked by increased incidence of senescent cell morphology with clusters of viable cells (Fig. 2B). After the spontaneous immortalization crisis, there were notable differences between the mackerel cells before and after this process. Morphologically, the cells became shorter and less variable in morphology (Fig. 2A, C). These morphological changes were confirmed with measurements of cell diameters before and after the crisis; 18.3 μm (± 0.53 SD) and 16.8 μm (± 0.17 SD), respectively (Fig. 2E). Importantly, cell doubling time also decreased significantly after the crisis, from 63.9 hr (± 19.1 SD) to 24.3 hr (± 4.91 SD) (Fig. 2D). During culture, the cells consistently tested negative for mycoplasma (Fig. S1). Additionally, early passage Mack2 cells (i.e., cells from the second isolation), demonstrated similar growth rates to that of the post-crisis Mack1 cells, with a doubling time of 22.9 hr (± 3.99 SD) (Fig. S2B).

**Figure 2:**
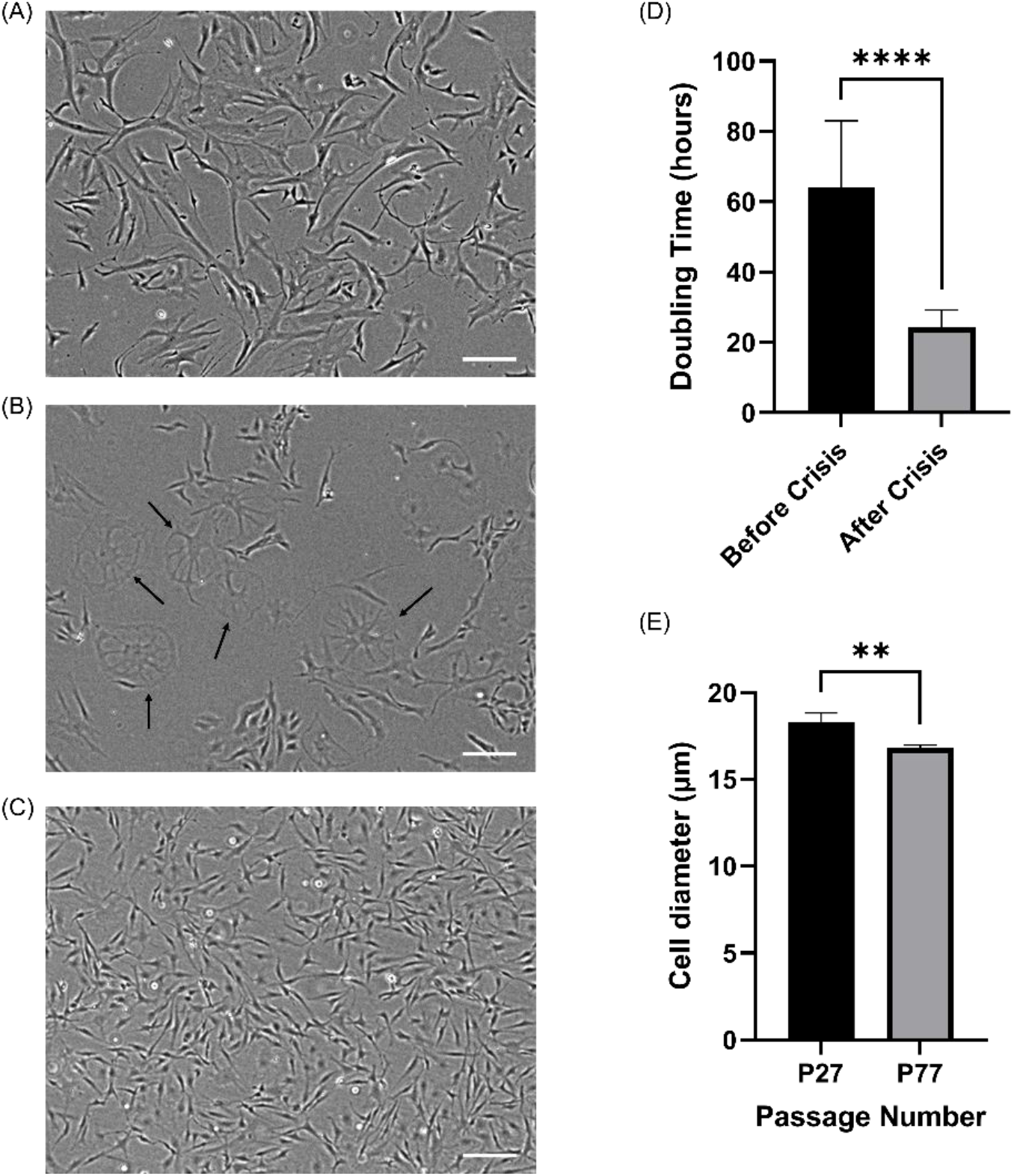
Spontaneous immortalization crisis event of Mack1 cells. (A-C) The progression of Mack1 cells through phase contrast imaging. Scale bars 200 μm. (A) Passage number 15, (B) passage number 42, (C) passage Number 84. Arrows in (B) indicate cells in crisis, as morphology is indicative of senescence. (D) Doubling time comparison between cells before crisis (passage numbers 12-26) and after crisis (passage numbers 114-128). Error bars indicate standard deviation, n = 15 distinct samples; statistical significance calculated by unpaired t-test and indicated by asterisks, in which p < 0.0001 (****). (E) Cell diameter comparison between cells before crisis (passage number 27) and after crisis (passage number 77). Error bars indicate standard deviation, n = 3 distinct samples; statistical significance calculated by unpaired t-test and indicated by asterisks, in which p < 0.01 (**).

### Confirmation of myogenicity

To confirm myogenicity, cells before and after crisis were continuously analyzed for PAX7 protein expression through immunocytochemistry. PAX7 is a specific marker of satellite cells and is detected through nuclear staining22,23. Cells before and after crisis stained for PAX7 (Fig. 3A), and image analysis showed a high percentage of PAX7+ nuclei, with cells after crisis nearing a pure satellite cell population (Fig. 3C). To confirm the cells potential to differentiate, cell feeding was ceased at 100% confluency, and, after 10 days of serum starvation, cells were immunostained for myosin heavy chain (MHC) with MF-20 (Fig. 3B). MF-20 staining confirmed cell differentiation into muscle fibers, with similar muscle morphology before and after the crisis. Despite cells before crisis being larger in diameter, image analysis showed no significant difference in muscle fiber diameter before and after the crisis (Fig. 3D). Mack2 cells were also tested for their differentiation potential and demonstrated functionality with robust muscle formation (Figs. S2C, D).

**Figure 3:**
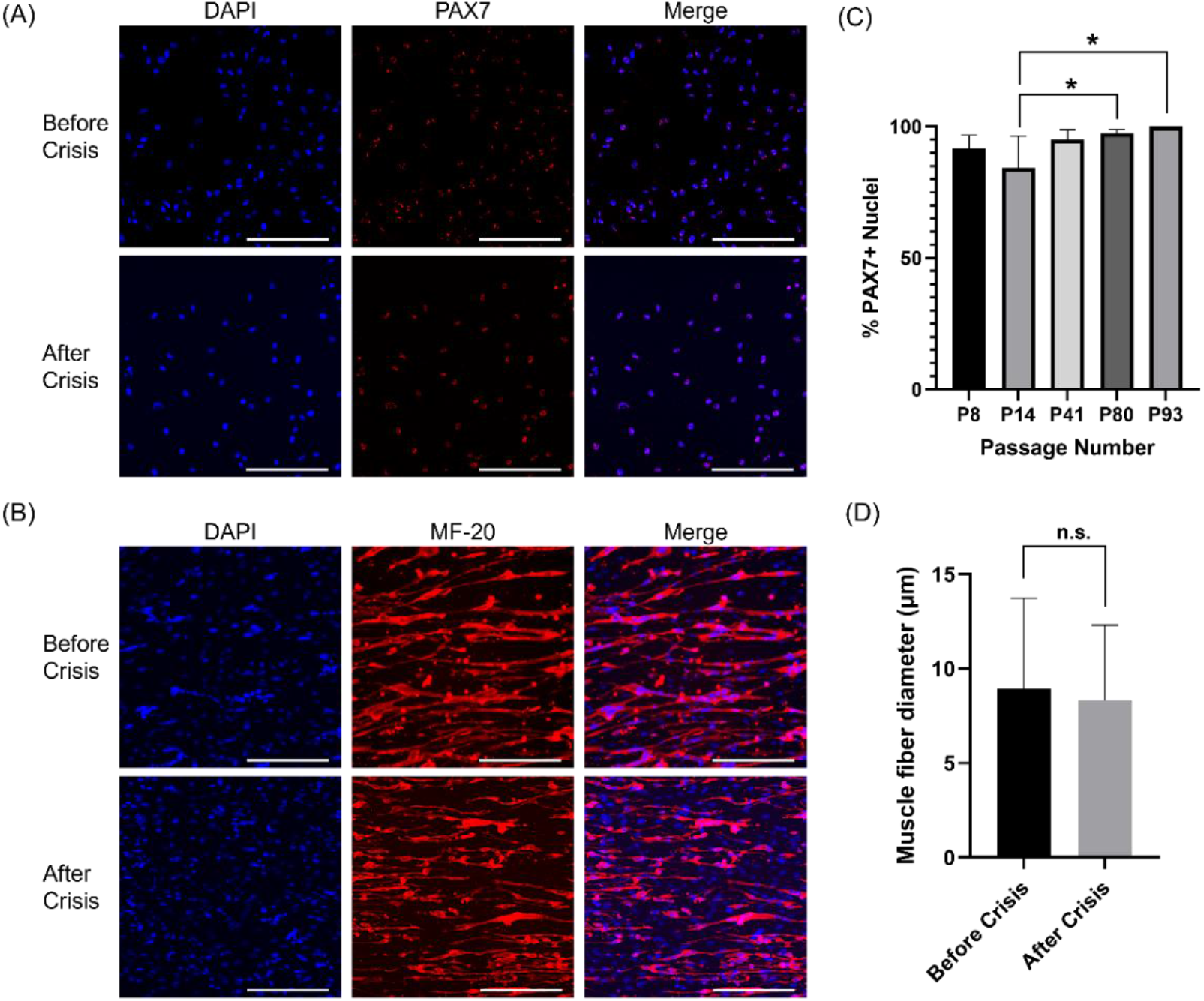
Myogenic potential and differentiation of Mack1 cells. (A) Immunofluorescence staining for cell nuclei (DAPI, blue) and PAX7 (red). Top and bottom panels are for passage numbers 14 and 93, respectively. Scale bars 100 μm. (B) Immunofluorescence staining for cell nuclei (DAPI, blue) and MHC (MF-20, red). Top and bottom panels are for passage numbers 8 and 56, respectively. Scale bars 100 μm. (C) Percent PAX7^+^ nuclei over multiple passages, as calculated through image analysis. Error bars indicate standard deviation, n = 3 or 6; statistical significance calculated by two-way ANOVA with multiple comparisons between the passage numbers and indicated by asterisks, in which p < 0.05 (*). (D) Muscle fiber diameter comparison between cells from passage numbers 8 and 93, through image analysis. Error bars indicate standard deviation, n = 20; no statistical significance calculated by unpaired t-test, in which n.s. is not significant.

### Lipid accumulation towards adipogenic-like phenotype

While muscle is the predominant component of fish fillets, fat plays an important role in flavor and texture. Fat is especially pronounced in mackerel, as it is often described as an oily fish^24,25^. Thus, to investigate the potential for mackerel muscle cells to accumulate lipids, induction towards an adipogenic-like phenotype was explored through media supplementation. Specifically, medium was supplemented with a lipid mixture and PPARG agonists (dexamethasone, 3-isobutyl-1-methylxanthine (IBMX), and insulin), as PPARG is the master regulator of adipogenic differentiation^26^. Inclusion of the lipid mixture with the agonists promoted lipid accumulation, confirmed through ORO staining and quantification of extracted ORO (Fig. 4A, B).

**Figure 4:**
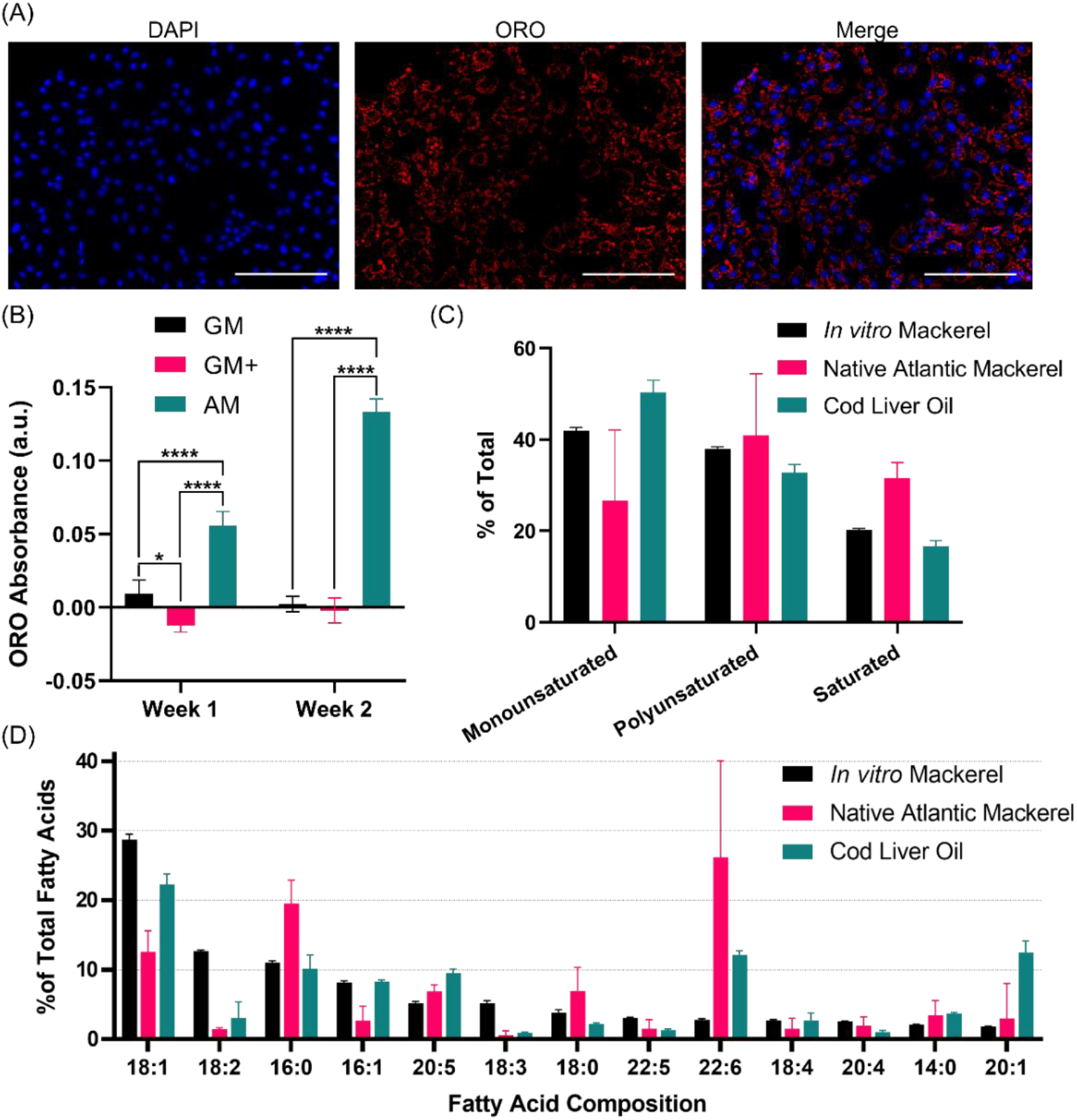
Lipid accumulation in Mack muscle cells. (A) Immunofluorescence staining for cell nuclei (DAPI, blue) and neutral lipids (Oil Red O, red). (B) Quantification of neutral lipid accumulation in different media through Oil Red O absorbance. GM: growth medium, GM+: growth medium + PPARG agonists, AM: adipogenic medium (GM+ with lipid mixture). (C) Percent of fatty acid saturation in adipogenic-like mackerel muscle cells, native Atlantic mackerel (adapted from ^25,27–29^), and cod liver oil (adapted from ^30,31^). Error bars indicate standard deviation, n = 3. (D) Fatty acid composition of adipogenic-like mackerel cells, native Atlantic mackerel (adapted from ^24,25,28,29^), and cod liver oil (adapted from ^30–32^). The top 13 most prevalent fatty acids from the *in vitro* mackerel are shown. Error bars indicate standard deviation, n = 3.

Lipidomics was used to characterize the fatty acid profiles of adipogenic-like cells and compared against published data for native Atlantic mackerel and cod liver oil, which is in the lipid mixture of the adipogenic medium. Fatty acid composition was binned into saturated fatty acids (SFAs), monounsaturated fatty acids (MUFAs), and polyunsaturated fatty acids (PUFAs) to compare lipid composition (Fig. 4C). In comparison to native mackerel, cultured cells contained relatively higher MUFA levels and lower SFA levels. In comparison to cod liver oil, the cultured cells contained lower levels of MUFAs. Specific fatty acid composition from the top 13 most prevalent fatty acids in the *in vitro* mackerel are exhibited in Fig. 4D. Of note, oleic (18:1) and linoleic (18:2) acids appear enriched in the *in vitro* samples, while docosahexaenoic acid (22:6) was lower (Fig. 4D). More specific investigation into the composition of the phospholipids, which constitute the cell membrane and are associated with taste, and triacylglycerides, which are the storage form of fat and constitute most of the lipids present, are presented in Fig. S4.

### Cell culture optimization

To understand the best practices for culturing mackerel cells, we explored seeding density, muscle differentiation regimes, and surface coatings. For seeding density, cells were seeded at 1,000 to 10,000 cell/cm^2^ to determine minimum seeding density and cell density at confluency. While cells seeded at 2,000 cells/cm^2^ proliferated well to roughly 30,000 cells/cm^2^ after 5 days, those at 1,000 cells/cm^2^ did not proliferate well after 5 days, indicating a minimum seeding density between 1,000 and 2,000 cells/cm^2^ (Fig. 5A). The maximum cell density (i.e., the cell density at 100% confluency) was determined to be ^~^45,000 cells/cm^2^, as by day 5 the cells seeded at densities of at least 4,000 cells/cm^2^ experienced contact inhibition and did not exceed ^~^45,000 cells/cm^2^ (Fig. 5A).

**Figure 5:**
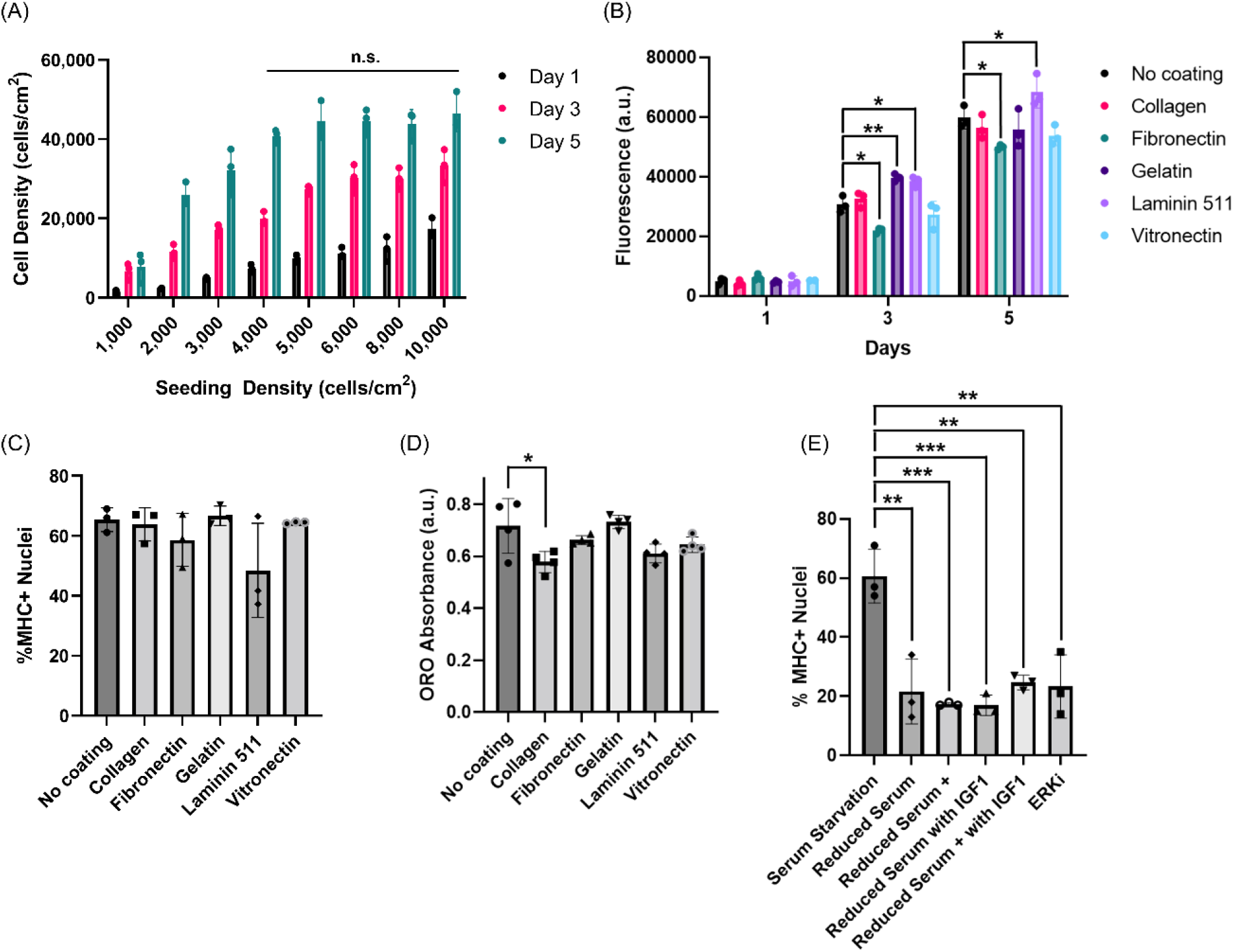
Cell culture optimization. (A) Comparative growth curves for varying seeding densities over days 1, 3, and 5. For day 5, two-way ANOVA indicated no significant differences between 4,000, 5,000, 6,000, 8,000, and 10,000 cell/cm^2^. (B) dsDNA quantification for Mack1 cells cultured on different ECM surface mimetics. Error bars indicate standard deviation, n =3; statistical significance calculated by two-way ANOVA with multiple comparisons between the surface mimetics and the no coating control at each time point and indicated by asterisks, in which p < 0.05 (*) and p < 0.01 (**). (C) Percent MHC+ nuclei for Mack1 cells cultured on different ECM surface mimetics. Error bars indicate standard deviation, n =3; statistical significance calculated by two-way ANOVA with multiple comparisons between the surface mimetics and the no coating control and indicated no significant differences between the mimetics. (D) Neutral lipid quantification, through Oil Red O extraction, for Mack1 cells cultured on different ECM surface mimetics. Error bars indicate standard deviation, n = 4; statistical significance calculated by two-way ANOVA with multiple comparisons between the surface mimetics and the no coating control and indicated by asterisks, in which p < 0.05 (*). (E) Percent MHC+ nuclei comparison between different muscle differentiation methods. Error bars indicate standard deviation, n =3; statistical significance calculated by two-way ANOVA with multiple comparisons between differentiation methods and indicated by asterisks, in which p < 0.01 (**) and p < 0.001(***).

Preferences for extracellular matrix (ECM) mimetics was investigated for more optimized growth, lipid accumulation, and muscle differentiation. For growth, laminin-511-E8 demonstrated increased proliferation rates (Fig. 5B), while for differentiation, the ECM mimetics did not improve differentiation efficiency of adipogenic-like differentiation or muscle differentiation (Fig. 5C, D).

Optimization for muscle differentiation involved investigation of differentiation regimes chosen based on previous literature approaches that demonstrated efficacy in mammalian cell differentiation^2,3,33^. The regimes involved reduction of serum to 2% in the differentiation medium, addition of transferrin, lysophosphatidic acid (LPA), and insulin (termed “reduced-serum +”), inclusion of ERK inhibitor (ERKi), and supplementation with IGF-1. When compared against the control of cessation of feeding (termed “serum starvation”), all of the tested regimes had significantly lower percentages of MHC+ cells (Fig. 5E).

### Gene expression

To build a research toolkit for the mackerel muscle cell line, a gene expression assay with RT-qPCR was developed. Due to the lack of availability of an annotated genome for *S. scombrus*, a reference genome for southern bluefin tuna (*Thunnus maccoyii*, NBCI RefSeq GCF_910596095.1) was originally used to design primers for the genes of interest: *HPRT* (hypoxanthine-guanine phosphoribosyltransferase, a housekeeping gene), *PAX3B* (paired-box 3b), *MYOD1* (myogenic differentiation 1), *MYOG* (myogenin), *TNNT3A* (troponin T type 3a), and *PPARG* (peroxisome proliferator-activated receptor gamma). The *T. maccoyii* genome was chosen because it is a fully annotated genome with genes of interest and because T. maccoyii is closely related to *S. scombrus^34^. After* initial screening for primer efficacy and specificity by running PCR products on DNA agarose gels, optimization was needed, due to either a lack of PCR product at the expected size or non-specific binding leading to multiple PCR products. To better investigate *S. scombrus* sequences, an Atlantic mackerel transcriptome (NCBI SRX2255766) was successfully aligned to the tuna genome, with 78% of reads uniquely mapped via the STAR aligner^35^. Of note, the 78% read mapping falls within the 70-90% observed when mapping human RNA Sequencing reads to a human reference genome^35^. Later iterations of primer design then referenced the *S. scombrus* annotated sequences. After successful confirmation where PCR products were singular and matched the expected size on agarose gels, the primer specificity was further confirmed through melt curves and Sanger sequencing (data not shown). Using the successful primers (Table S1), gene expression during proliferation, muscle differentiation through serum starvation (cessation of feeding), muscle differentiation through serum reduction (2% FBS) with additives, and lipid accumulation was investigated (Figs. 6, S4). Interestingly, the adipogenic-like cells generally followed trends similar to muscle differentiation and did not exhibit increases in *PPARG. PAX3B* is a marker of satellite cell stemness, while *MYOD1, MYOG*, and *TNNT3A* are all genes associated with muscle differentiation. Thus, the decrease of *PAX3B* and increases of *MYOD1, MYOG*, and *TNNT3A* during differentiation were expected.

**Figure 6:**
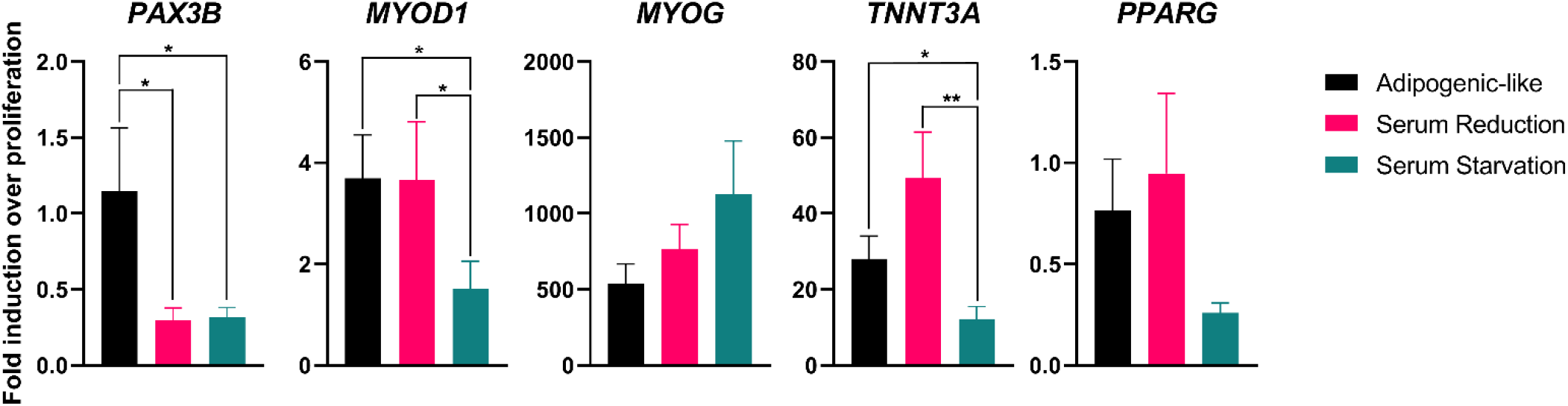
RT-qPCR Gene Expression for PAX3B, MYOD, MYOG, TNNT3A, and PPARG. Fold induction over proliferation is represented as 2^-ΔΔCt^. Error bars indicate standard error of the mean, n = 9 (n = 3 experimental, n = 3 technical, outliers removed); statistical significance calculated by two-way ANOVA with multiple comparisons between the differentiation samples and indicated by asterisks, in which p < 0.05 (*) and p < 0.01 (**).

## Discussion

Cellular aquaculture offers many potential advantages over conventional fishing and aquaculture. However, the lack of fish muscle cell lines has hindered cultured meat research for fish, such as studies into serum-free media, 3D expansion options, and structured tissue formation. While most cellular agriculture research involves the use of bovine satellite cells due to their food and environmental relevance, traditional animal muscle cell and tissue *in vitro* studies have focused on the mouse myoblast C2C12 cell line due to availability^36,37^. The present study provides a new option for the field of fish cell and tissue culture, through the establishment of a mackerel skeletal muscle cell line.

The Atlantic mackerel cell line Mack1 is stable, having progressed out of a spontaneous immortalization crisis event and proliferating indefinitely since the crisis (Fig. 2). The spontaneous immortalization method of establishing a cell line is preferred over genetic modification methods, as genetically modified organisms (GMOs) will face stricter regulation, especially in Europe^38^. Mack1 is a highly pure muscle cell line, confirmed through immunostaining for PAX7 and MHC for myogenic potential and differentiation, respectively (Fig. 3). Mack1 cells also possess the ability to accumulate lipids, demonstrating potential for both fat and muscle phenotypes (Fig. 4). Optimization of Mack1 cell culture and research tool development was then explored through seeding density and differentiation studies, as well as RT-qPCR assay development (Figs. 5, 6).

However, further understanding and optimization of cell culture are needed. While the expansion of the cells resulted in doubling times of 24.3 hr (± 4.91 SD), serum-free medium without FBS is necessary to reduce costs and to address ethical considerations and animal-free ingredients in cellular agriculture. Methods that work well with mammalian cell culture are not interchangeable with fish cell culture, as evaluations with Beefy9 serum-free medium (data not shown)^39^ and ERK inhibition (Fig. 5B)^2^ were unsuccessful. To develop a serum-free medium for fish cell culture, media with mushroom extracts and insulin-like growth factor 2 (IGF-2) have shown promise in goldfish and medaka (*Oryzias latipes*) cultures, respectively^40,41^. However, neither medium promoted proliferation of Mack1 cells (data not shown). Additionally, while the Mack1 cells did not demonstrate a reliance on an ECM mimetic for cultivation (Fig. 5B), the surface coating screens should be revisited during serum-free media development, as the FBS includes proteins that likely adsorb onto tissue culture plastic and encourage cell adhesion.

While demonstration of myogenic and adipogenic-like phenotypes underscore cell functionality, the continued development of research tools to understand biological and molecular processes should accelerate progress towards enhanced functional utility for these cells. While the Mack cells stained positively for PAX7 and MHC, staining with antibodies for MYOD and MYOG was unsuccessful (data not shown). The unsuccessful trials highlight the need for future research tool development, to both pose and answer questions regarding cell line function and development. For example, the RT-qPCR assay introduced questions regarding temporal gene and protein expression patterns in muscle differentiation, while the lack of *PPARG* gene expression in lipid accumulation indicated the adipogenic-like cells likely undergo phenotypic changes rather than a proper transdifferentiation. For muscle differentiation, despite a lower percentage of MHC+ nuclei, the reduced serum differentiation regime resulted in increased gene expression of *TNNT3A* when compared to serum starvation conditions (Figs. 5B, 6). While the gene and protein in the assays are different, TNNT3A and MHC are interrelated, as the troponin T bridges myosin heads to tropomyosin in differentiated muscle^43,44^. The apparent discrepancy between *TNNT3A* gene expression and MHC protein expression could indicate a temporal difference in protein and gene expression or a difference in troponin T to MHC ratios during stages of differentiation. While mackerel muscle differentiation regimes generally followed expected gene expression patterns (i.e., a decrease in *PAX7* and increases in *MYOD, MYOG*, and *TNNT3A*), lipid accumulation unexpectedly followed similar patterns, without a change in *PPARG* (Fig. 6). Previous studies with transdifferentiated C2C12s also noted the lack of expression of *PPARG* in adipocyte-like cells; instead, increases in adipocyte lipid binding protein (*ALBP*), cyclooxygenase-2 (*COX2*), and fatty acid-activated receptor (*FAAR*) were observed^45,46^. Because the Mack1 cells are muscle cells, it is likely that cell-to-cell signaling/contact initiated intercellular communication for muscle differentiation^47^. Nonetheless, the specific mechanism of lipid accumulation in the Mack muscle cells is elusive. When C2C12s were fed with fatty acids and thiazolidinediones, *MYOG* expression was halted, although for the Mack cells *MYOG* expression increased during lipid accumulation^46^. Despite gaps in understanding the mechanisms of these phenotypic changes, observing increased lipid accumulation was promising as a fat analog for cultivated foods.

Lipid accumulation in adipogenic medium was significant in comparison to growth medium (Fig. 3B), allowing for investigation into the lipid profile via lipidomics (Fig. 3C, D and Fig. S4). Comparing lipid composition generated with these cells to native mackerel is limited due to the large variation in native mackerel lipid content seasonally and geographically^25^. However, an interesting observation was the relatively high levels of saturated fatty acids in the phospholipid fraction (Fig. S4C). A hypothesis is that since the fish cells are cultured at 27°C, a temperature higher than that of the Atlantic Ocean where the fish was caught, the phospholipid bilayer of the cells may be adapted to higher saturation levels to maintain solidified states. *In vivo* studies have found that fish from warmer climates generally have higher saturated fat phospholipid fractions than fish from cold water climates, with *T. bernacchii* increasing saturated fat levels when acclimated to a warmer water temperature^48,49^.

The present work details the establishment and characterization of a fish muscle cell line that could serve as a characterization framework for other cellular agriculture cell line developments. The successful isolation and propagation of the Mack cells were achieved only after multiple isolation failures for different species and with different protocols. Some observations from the attempted isolations were the need for robust sterilization methods for the tissue, varying efficacy of enzymatic agents (e.g., trypsin, collagenase, Accumax) for different species, and the need for debris removal through pre-plate methods (i.e., allowing cells and debris adhesion to an initial cell culture vessel and then moving the non-adherent material to a new vessel) and subsequent washes after cell adhesion. While we hypothesized freshness and the time between the death of the fish and the cell isolation would be key parameters in success, surprisingly, that was not the case for the two mackerel isolations. The fish were transported as whole fish on ice, and the isolations were not performed until about 5 hr *post-mortem*. Despite the time between fish death and cell isolation, the two mackerel cell isolations were more successful in comparison to cell isolations from other species (salmon, zebrafish, tilapia, trout, tuna), both in terms of yield of isolated cells per gram of tissue and in growth rate of the cells after isolation.

For future research, removal of all animal components, notably FBS in the growth medium and cod liver oil in the adipogenic medium, will be key to the future utility of these cells in cellular agriculture. To replace the cod liver oil, plant-based alternatives including intralipid or purified γ-linolenic acid from borage seed oil can be pursued^21,50,51^. For optimization of cell culture conditions and removal of animal components, further understanding of the biological and molecular mechanisms of the cells should accelerate further progress. However, since *S. scombrus* and fish in general are relatively understudied from an *in vitro* cell culture perspective, understanding cell processes are more challenging due to the lack of suitable experimental tools. In developing RT-qPCR, we circumvented the lack of an annotated genome by aligning a mackerel transcriptome with an annotated genome from southern bluefin tuna, a closely related species. This sequence similarity between the species was high^34^, and future studies on species without annotated genomes could also leverage phylogeny in determining potential reference genomes. While RT-qPCR is one step in developing research tools, omics techniques should provide important insight by gathering large datasets to infer gene, metabolite, or protein functions. Transcriptomics and proteomics will inform sequences of genes and proteins specific to Atlantic mackerel, which could further inform the design of primers and antibodies to study the cells with more resolution. Optimization of cell characterization will enhance cell culture and accelerate development of tissue cultures from these cells. Ultimately, a proof-of-concept 3D tissue construct, with both muscle and fat generation, will be invaluable in showcasing the potential value of the Mack cells for industrial applications.

## Acknowledgements

We thank the Good Food Institute, the United States Department of Agriculture (2021-69012-35978), and Wanda Fish Technologies for support of this work. We thank Jake Marko, Sophie Letcher, Andrew Stout, Natalie Rubio, Rebecca Batorsky, Christel Andreassen, Edward Gordon, and Claire Bomkamp for their scientific input. We also thank Captains Johnny Johnson, Ira Shank, Pete Speeches, and Andrew Lebel for their graciousness in providing fish tissue for this research.

## Cell Line Availability

The spontaneously immortalized MACK1 cells are available for non-profit researchers through Kerafast (Catalog # ETU008-FP).

## Notes

### Competing Interest Statement

The authors have declared no competing interest.

## References

1. Stout AJ, Mirliani AB, Soule-albridge EL, Cohen JM, Kaplan L. Engineering carotenoid production in mammalian cells for nutritionally enhanced cell-cultured foods. Metab Eng. 2020;62(July):126–137. doi:10.1016/j.ymben.2020.07.011

2. Eigler T, Zarfati G, Amzallag E, et al. ERK1/2 inhibition promotes robust myotube growth via CaMKII activation resulting in myoblast-to-myotube fusion. Dev Cell. 2021;56(24):3349–3363.e6. doi:10.1016/j.devcel.2021.11.022

3. Messmer T, Klevernic I, Furquim C, et al. A serum-free media formulation for cultured meat production supports bovine satellite cell differentiation in the absence of serum starvation. Nat Food. 2022;3:74–85. doi:10.1038/s43016-021-00419-1

4. Ben-Arye T, Shandalov Y, Ben-Shaul S, et al. Textured soy protein scaffolds enable the generation of three-dimensional bovine skeletal muscle tissue for cell-based meat. Nat Food. 2020;1(4):210–220. doi:10.1038/s43016-020-0046-5

5. Béné C, Barange M, Subasinghe R. Feeding 9 billion by 2050 – Putting fish back on the menu. Food Secur. Published online 2015:261–274. doi:10.1007/s12571-015-0427-z

6. Myers RA, Worm B. Rapid worldwide depletion of predatory fish communities. Nature. 2003;423(May):280–283.

7. Funk CC, Brown ME. Declining global per capita agricultural production and warming oceans threaten food security. Food Secur. 2009;(October 2008):271–289. doi:10.1007/s12571-009-0026-y

8. Walker PJ, Winton JR. Emerging viral diseases of fish and shrimp. Vet Res. 2010;41(6). doi:10.1051/vetres/2010022

9. Rubio N, Datar I, Stachura D, Kaplan D, Krueger K. Cell-Based Fish: A Novel Approach to Seafood Production and an Opportunity for Cellular Agriculture. Front Sustain Food Syst. 2019;3. doi:10.3389/fsufs.2019.00043

10. Bairoch A. The cellosaurus, a cell-line knowledge resource. J Biomol Tech. 2018;29(2):25–38. doi:10.7171/jbt.18-2902-002

11. Vishnolia KK, Martin NRW, Player DJ, Spikings E, Lewis MP. Zebrafish skeletal muscle cell cultures: Monolayer to three-dimensional tissue engineered collagen constructs. bioRxiv. 2020;44(0). doi:10.1101/2020.12.10.419168

12. Li N, Guo L, Guo H. Establishment, characterization, and transfection potential of a new continuous fish cell line (CAM) derived from the muscle tissue of grass goldfish (Carassius auratus). Vitr Cell Dev Biol - Anim. Published online 2021:912–931. doi:10.1007/s11626-021-00622-1

13. Gabillard JC, Sabin N, Paboeuf G. In vitro characterization of proliferation and differentiation of trout satellite cells. Cell Tissue Res. Published online 2010:471–477. doi:10.1007/s00441-010-1071-8

14. Oestbye TK, Ytteborg E. Preparation and Culturing of Atlantic Salmon Muscle Cells for In Vitro Studies. In: Myogenesis: Methods and Protocols. Vol 1889.; 2019:319–330. doi:10.1007/978-1-4939-8897-6_19

15. Specht-Overholt S, Romans JR, Marchello MJ, et al. Fatty Acid Composition of Commercially Manufactured Omega-3 Enriched Pork Products, Haddock, and Mackerel. J Anim Sci. 1997;75(9):2335–2343.

16. Monterey Bay Aquarium. Seafood Watch - Atlantic Mackerel. Published 2022. https://www.seafoodwatch.org/recommendation/mackerel/atlantic-mackerel-15404?species=315

17. FDA/EPA. FDA/EPA 2004 Advice on What You Need to Know About Mercury in Fish and Shellfish. Published 2004. https://www.fda.gov/food/metals-and-your-food/fdaepa-2004-advice-what-you-need-know-about-mercury-fish-and-shellfish

18. Ward RD, Hanner R, Hebert PDN. The campaign to DNA barcode all fishes, FISH-BOL. J Fish Biol. 2009;74:329–356. doi:10.1111/j.1095-8649.2008.02080.x

19. Aranishi F. PCR-RFLP Analysis of Nuclear Nontranscribed Spacer for Mackerel Species Identification. J Agric Food Chem. Published online 2005:508–511.

20. Breitkopf SB, Yuan M, Xu Y, Peake DA, Manning BD, Asara JM. A Relative Quantitative Positive/Negative Ion Switching Method for Untargeted Lipidomics via High Resolution LC-MS/MS from Any Biological Source. Vol 13.; 2018. doi:10.1007/s11306-016-1157-8

21. Yuen Jr. JS, Saad MK, Xiang N, Barrick BM, Dicindio H, Li C. Macroscale Adipose Tissue from Cellular Aggregates: A Simplified Method of Mass Producing Cell-Cultured Fat for Food Applications. bioRxiv. Published online 2022.

22. Relaix F, Montarras D, Zaffran S, et al. Pax3 and Pax7 have distinct and overlapping functions in adult muscle progenitor cells. J Cell Biol. 2006;172(1):91–102. doi:10.1083/jcb.200508044

23. Seale P, Sabourin LA, Girgis-gabardo A, Mansouri A, Gruss P, Rudnicki MA. Pax7 Is Required for the Specification of Myogenic Satellite Cells. Cell. 2000;102:777–786.

24. Aminullah AKM, Ratnayake WMN, Ackman RG. Effect of Smoking on the Proximate Composition Atlantic Mackerel (Scomber scombrus). J Food Sci. 1986;51(2).

25. Oudiani S El, Chetoui I, Darej C, Moujahed N. Sex and seasonal variation in proximate composition and fatty acid profile of Scomber scombrus (L. 1758) fillets from the Middle East Coast of Tunisia. Grasas y aceites. 2019;70(1):1–10.

26. Ahmadian M, Suh JM, Hah N, et al. PPARγ signaling and metabolism: the good, the bad and the future. Nat Med. 2013;19(5). doi:10.1038/nm.3159.PPAR

27. Boselli E, Pacetti D, Lucci P, Frega NG. Characterization of Phospholipid Molecular Species in the Edible Parts of Bony Fish and Shellfish. J Agric Food Chem. 2012;60:3234–3245.

28. Bhuiyan AKMA, Ratnayake WMN, Ackman RG. Stability of Lipids and Polyunsaturated Fatty Acids During Smoking of Atlantic Mackerel (Scomber scombrus L.). J Am Oil Chem Soc. 1985;63(3):324–328.

29. Molversmyr E. Identification and quantitation of lipids in Atlantic mackerel (Scomber scombrus), wild and farmed Atlantic salmon (Salmo salar), and salmon feed by GC-MS. Published online 2020.

30. Lamberstein G, Braekkan OR. The Fatty Acid Composition of Cod Liver Oil. Reports Technol Res Concern Nor Fish Ind. 1965;IV(11).

31. Helland I, Saarem K, Saugstad O, Drevon C. Fatty acid composition in maternal milk and plasma during supplementation with cod liver oil. Eur J Clin Nutr. 1998;52:839–845. doi:10.1038/sj.ejcn.1600656

32. Pan SX, Ushio H, Ohshima T. Comparison of volatile compounds formed by autoxidation and photosensitized oxidation of cod liver oil in emulsion systems. Fish Sci. 2005;71(June):639–647. doi:10.1111/j.1444-2906.2005.01010.x

33. Furuhashi M, Morimoto Y, Shima A, Nakamura F, Ishikawa H, Takeuchi S. Formation of contractile 3D bovine muscle tissue for construction of millimetre-thick cultured steak. npj Sci Food. Published online 2021:1–8. doi:10.1038/s41538-021-00090-7

34. Miya M, Friedman M, Satoh TP, et al. Evolutionary Origin of the Scombridae (Tunas and Mackerels): Members of a Paleogene Adaptive Radiation with 14 Other Pelagic Fish Families. PLoS One. 2013;8(9):1–19. doi:10.1371/journal.pone.0073535

35. Dobin A, Davis CA, Schlesinger F, et al. STAR: ultrafast universal RNA-seq aligner. Bioinformatics. 2013;29(1):15–21. doi:10.1093/bioinformatics/bts635

36. Blau HM, Chiu C, Webster C. Cytoplasmic Activation of Human Nuclear Genes in Stable Heterocaryons. Cell. 1983;32(April):1171–1180.

37. Yaffe D, Saxel O. Serial passaging and differentiation of myogenic cells isolated from dystrophic mouse muscle. Nature. 1977;270(December):725–727.

38. Post MJ, Levenberg S, Kaplan DL, et al. Scientific, sustainability and regulatory challenges of cultured meat. Nat Food. 2020;1(7):403–415. doi:10.1038/s43016-020-0112-z

39. Stout AJ, Mirliani AB, Rittenberg ML, et al. Simple and effective serum-free medium for sustained expansion of bovine satellite cells for cell cultured meat. Commun Biol. 2022;5(1). doi:10.1038/s42003-022-03423-8

40. Benjaminson MA, Gilchriest JA, Lorenz M. In vitro edible muscle protein production system (MPPS): Stage 1, fish. Acta Astronaut. 2002;51(12):879–889. doi:10.1016/S0094-5765(02)00033-4

41. Yuan Y, Hong Y. Medaka insulin-like growth factor-2 supports self-renewal of the embryonic stem cell line and blastomeres in vitro. Sci Rep. 2017;7(1):1–11. doi:10.1038/s41598-017-00094-y

42. Yao T, Asayama Y. Animal-cell culture media: History, characteristics, and current issues. Reprod Med Biol. 2017;16(2):99–117. doi:10.1002/rmb2.12024

43. Rasmussen M, Jin J-P. Troponin Variants as Markers of Skeletal Muscle Health and Diseases. Front Physiol. 2021;12(September):1–14. doi:10.3389/fphys.2021.747214

44. Muroya S, Nakajima I, Oe M, Chikuni K. Effect of phase limited inhibition of MyoD expression on the terminal differentiation of bovine myoblasts: No alteration of Myf5 or myogenin expression. Dev Growth, Differ. 2005;47:483–492.

45. Chen X, Xu S, Wei S, Deng Y, Li Y, Yang F. Comparative Proteomic Study of Fatty Acid-treated Myoblasts Reveals Role of Cox-2 in Palmitate-induced Insulin Resistance. Sci Rep. 2016;6(February):1–12. doi:10.1038/srep21454

46. Teboul L, Gaillard D, Staccini L, Inadera H, Amri E, Grimaldi PA. Thiazolidinediones and Fatty Acids Convert Myogenic Cells into Adipose-like Cells. J Biol Chem. 1995;270(47):28183–28187. doi:10.1074/jbc.270.47.28183

47. Demonbreun AR, McNally EM. Muscle cell communication in development and repair. Curr Opin Pharmacol. 2018;34(June):1–15. doi:10.1016/j.coph.2017.03.008.Muscle

48. Patton JS. The effect of pressure and temperature on phospholipid and triglyceride fatty acids of fish white msucle: a comparison of deepwater and surface marine species. Comp Biochem Physiol. 1975;52B:105–110.

49. Malekar VC, Morton JD, Hider RN, Cruickshank RH, Hodge S, Metcalf VJ. Effect of elevated temperature on membrane lipid saturation in Antarctic notothenioid fish. PeerJ. 2018;6(e4765):1–25. doi:10.7717/peerj.4765

50. Starkey JD, Yamamoto M, Yamamoto S, Goldhamer DJ. Skeletal Muscle Satellite Cells Are Committed to Myogenesis and Do Not Spontaneously Adopt Nonmyogenic Fates. J Histochem Cytocehmistry. 2011;59(1):33–46. doi:10.1369/jhc.2010.956995

51. Wada MR, Inagawa-ogashiwa M, Shimizu S, Yasumoto S. Generation of different fates from multipotent muscle stem cells. Development. 2002;129:2987–2995.

