## Supplemental Tables and Figures for "Continuous Fish Muscle Cell Line with Capacity for Myogenic and Adipogenic-like Phenotypes"

### Supplementary Information

**Supplementary Table 1: Media Formulations**

| **Component** | **Concentration** |
| --- | --- |
| **L-15 Growth Medium** |  |
| Leibovitz’s L-15 Medium (ThermoFisher Scientific #11415064) |  |
| Fetal bovine serum (FBS, ThermoFisher Scientific #26140079) | 20% |
| Recombinant human fibroblast growth factor (FGF-basic 154 a.a.; PeproTech #100-18B) | 1 ng/mL |
| HEPES, pH 7.4 (Sigma Aldrich #H4034) | 20 mM |
| Antibiotic-Antimycotic (ThermoFisher Scientific #1540062) | 1% |
| Gentamicin solution (Sigma Aldrich #G1397) | 10 µg/mL |
| **Adipogenic Medium** |  |
| L-15 growth medium (described above) |  |
| Insulin solution from bovine pancreas (Sigma Aldrich #I0516) | 10 µg/mL |
| 3-isobutyl-1-methylxanthine (IBMX, Sigma Aldrich #I5879) | 0.5 mM |
| Dexamethasone (Sigma Aldrich #D4902) | 0.25 μM |
| Lipid mixture (Sigma Aldrich #L5146) | 10 µL/mL |
| **Reduced Serum Medium** |  |
| Leibovitz’s L-15 Medium (ThermoFisher Scientific #11415064) |  |
| Fetal bovine serum (FBS, ThermoFisher Scientific #26140079) | 2% |
| HEPES, pH 7.4 (Sigma Aldrich #H4034) | 20 mM |
| Antibiotic-Antimycotic (ThermoFisher Scientific #1540062) | 1% |
| Gentamicin solution (Sigma Aldrich #G1397) | 10 µg/mL |
| +/- insulin-like growth factor 1 (IGF-1, Shenandoah Biotechnology #100-34) | 100 ng/mL |
| **Reduced Serum + Additives** |  |
| Reduced serum medium (described above) |  |
| 1-Oleoyl Lysophosphatidic Acid (LPA, Fisher Scientific #NC9401387) | 10 µM |
| Optiferrin® recombinant human transferrin (InVitria #777TRF029) | 135 nM |
| Insulin solution from bovine pancreas (Sigma Aldrich #I0516) | 1.8 µM |
| +/- insulin-like growth factor 1 (IGF-1, Shenandoah Biotechnology #100-34) | 100 ng/mL |
| **ERKi Medium** |  |
| Reduced serum medium (described above) |  |
| Calbiochem™ ERK Inhibitor (ERKi, Sigma Aldrich #328006) | 2 µM |

**Supplementary Table 2: RT-qPCR Primers**

| **Gene** | **NCBI Accession Number** | **Primer Direction** | **Primer Sequence (5’ – 3’)** |
| --- | --- | --- | --- |
| *HPRT* | XM_042419962.1 | Fwd | GTCTACGTTGACAGGCAAGAATGT |
|  |  | Rev | GTCTGGTCGGTAGCCAACACT |
| *PAX3B* | XM_042415392.1 | Fwd | AAGCGGGAAAACCCGGTGAG |
|  |  | Rev | CCACATCCGAACCCTCGTCT |
| *MYOD1* | XM_042412988.1 | Fwd | TTGGAGCACTACAGCGGGGA |
|  |  | Rev | GCTGGTGTCGGTACTGATCCG |
| *MYOG* | XM_042406865.1 | Fwd | GGAGCACCCTGATGAACCCC |
|  |  | Rev | CGCTTGACGACGACACTCTGG |
| *TNNT3A* | XM_042412366.1 | Fwd | TCAGCGCGGTAAGTTTGCAG |
|  |  | Rev | CTCCTCTTCTACGGCCTCGACA |
| *PPARG* | XM_042402801.1 | Fwd | CCATGCCTGTGAGGGCTGTA |
|  |  | Rev | GGCCAAAACGAATGGCGTTG |

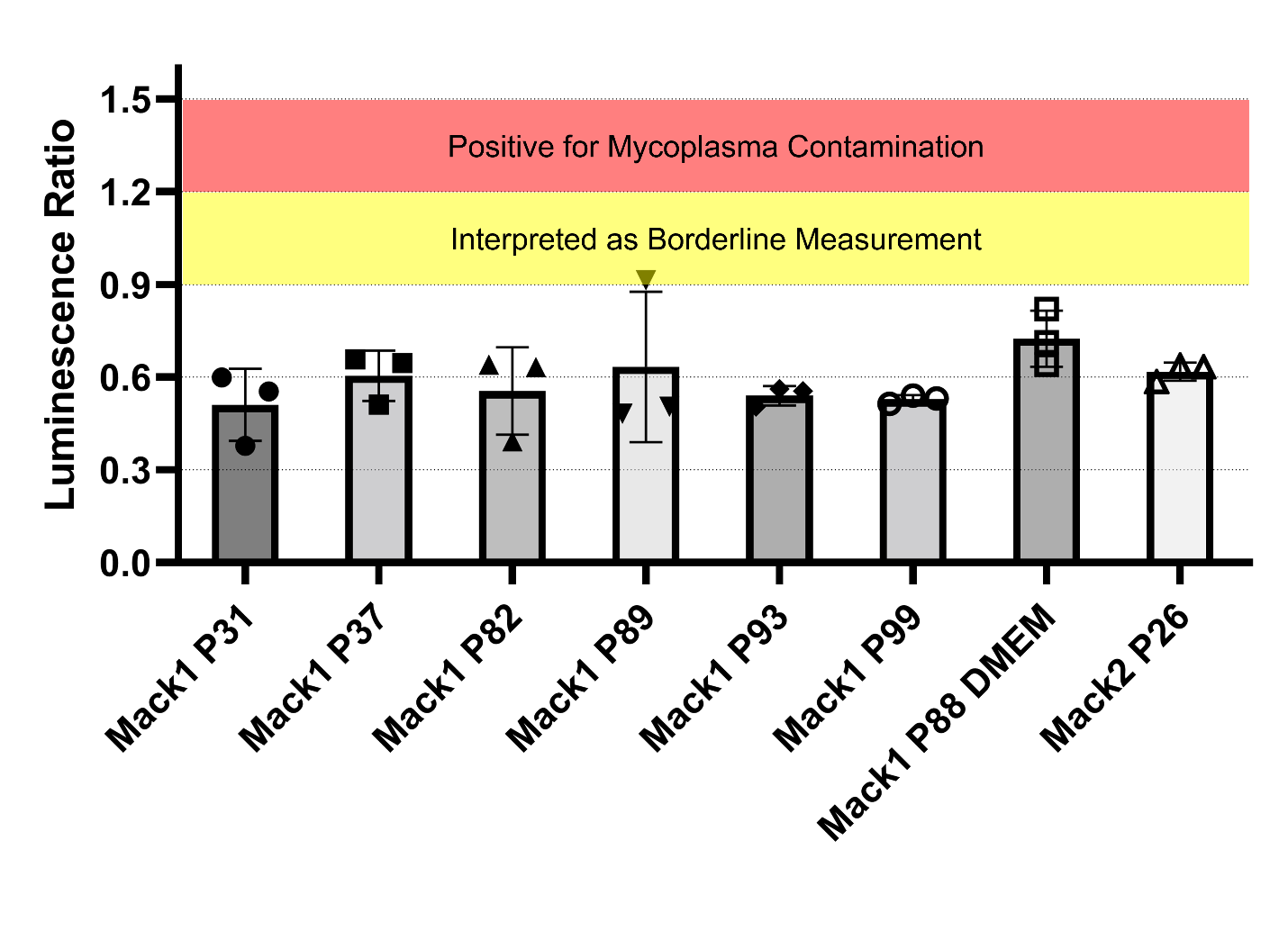

**Figure S1: Mycoplasma testing**, as measured by MycoAlert® Mycoplasma Detection Kit. Luminescence ratio values between 0.9 and 1.2 indicate samples may be borderline for contamination, while values above 1.2 indicate contamination. Error bars indicate standard deviation, n =3.

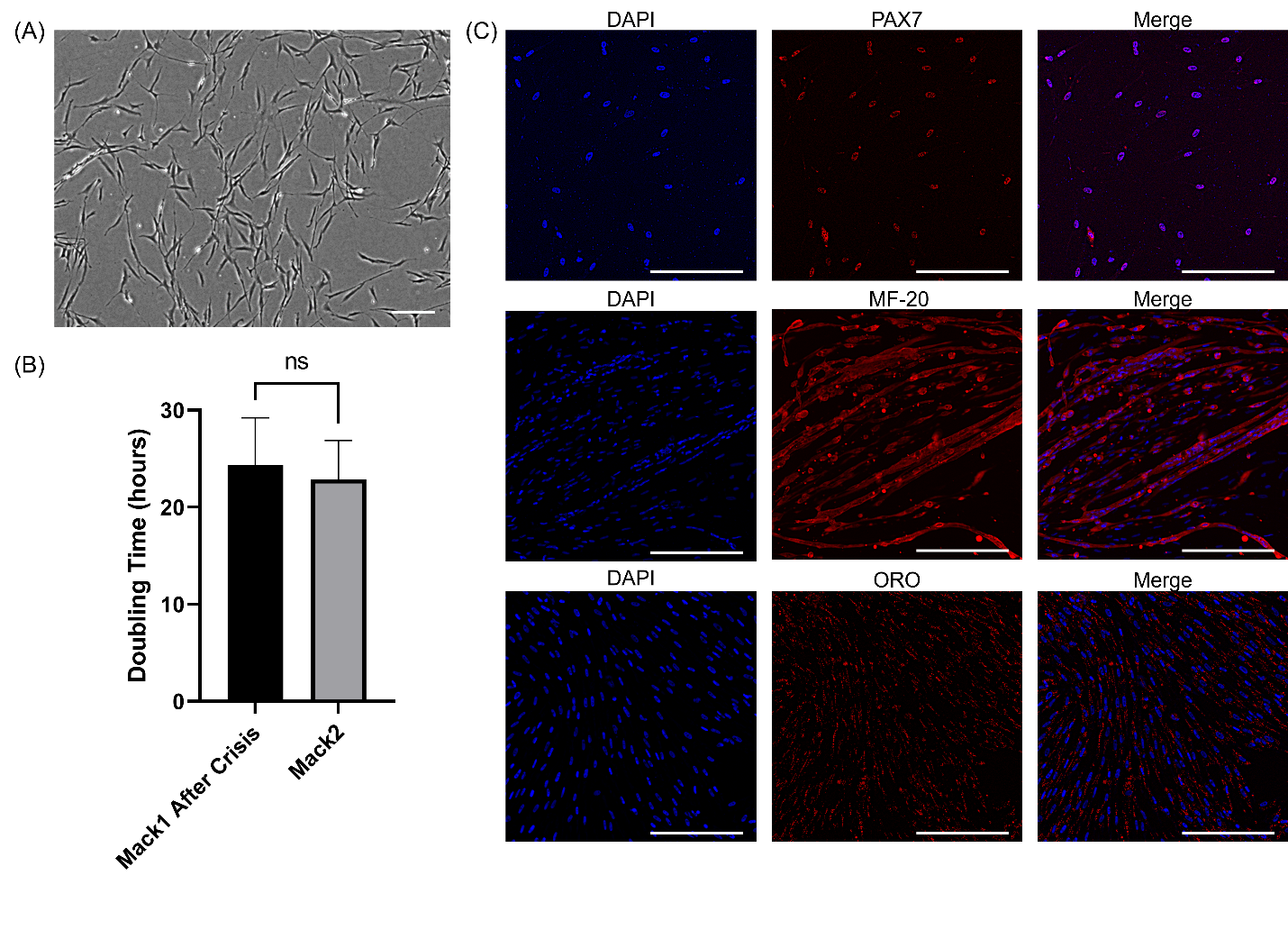

**Figure S2: Mack2 characterization.** (A) Phase contrast imaging of Mack2 P31. Scale bar 200 μm. (B) Doubling time comparison between Mack1 after crisis (passage numbers 114-128) and Mack2 (passage numbers 39-53). Error bars indicate standard deviation, n = 15; statistical significance calculated by unpaired t-test, in which n.s. is not significant. (C) Immunofluorescent staining on Mack2 P27 for cell nuclei (DAPI, blue), PAX7 (red, top panel), MHC (MF-20, red, middle panel), and Oil Red O (red, bottom panel) during proliferation, muscle differentiation, and adipogenic-like differentiation, respectively. Scale bars 100 μm.

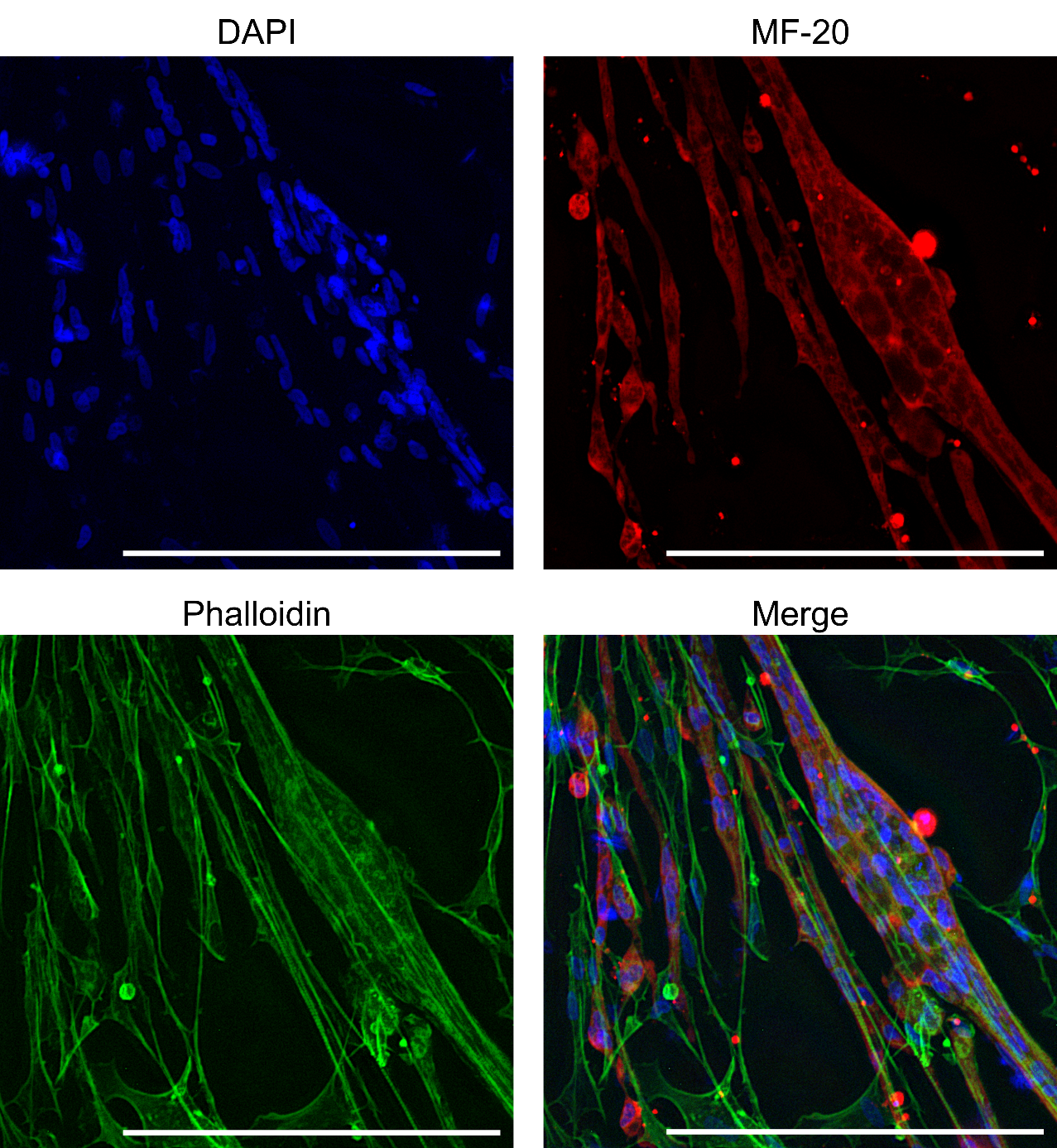

**Figure S3: Mack2 muscle differentiation**. Immunofluorescent staining for cell nuclei (DAPI, blue, top left), MHC (MF-20, red, top right), F-actin (Phalloidin, green, bottom left), and merge of all three (bottom right). Scale bars 200 μm.

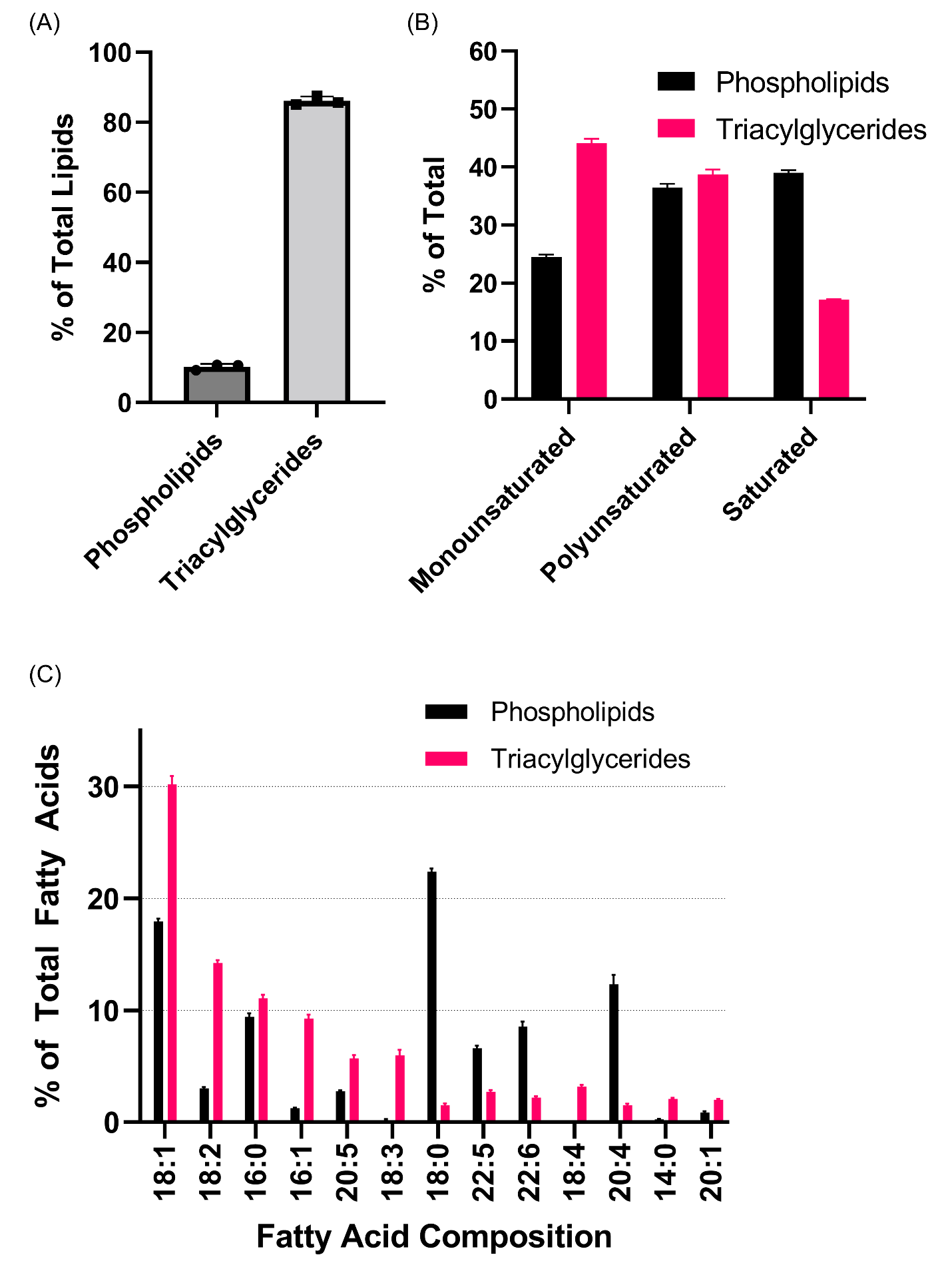

**Figure S4: Phospholipid (PL) and triaccylglyceride (TAG) lipidomics of adipogenic-like mackerel cells.** (A) The percentage of PL and TAG, respective to the total lipids detected in lipidomics. (B) Percent of fatty acid saturation in the PL and TAG fractions. (C) The top 13 most prevalent fatty acids from the in vitro mackerel PL and TAG fractions are shown. Error bars indicate standard deviation, n = 3.
